# Cooperation among *c*-subunits of F_o_F_1_-ATP synthase in rotation-coupled proton translocation

**DOI:** 10.1101/2021.03.25.436961

**Authors:** Noriyo Mitome, Shintaroh Kubo, Sumie Ohta, Hikaru Takashima, Yuto Shigefuji, Toru Niina, Shoji Takada

## Abstract

In F_o_F_1_-ATP synthase, proton translocation through F_o_ drives rotation of the *c*-subunit oligomeric ring relative to the *a*-subunit. Recent studies suggest that in each step of the rotation, key glutamic acid residues in different *c*-subunits contribute to proton release to and proton uptake from the *a*-subunit. However, no studies have demonstrated cooperativity among *c*-subunits toward F_o_F_1_-ATP synthase activity. Here, we addressed this using *Bacillus* PS3 ATP synthase harboring *c*-ring with various combinations of wild-type and *c*E56D, enabled by genetically fused single-chain *c*-ring. ATP synthesis and proton pump activities were decreased by a single *c*E56D mutation and further decreased by double *c*E56D mutations. Moreover, activity further decreased as the two mutation sites were separated, indicating cooperation among *c*-subunits. Similar results were obtained for proton transfer-coupled molecular simulations. Simulations revealed that prolonged proton uptake in mutated *c*-subunits is shared between two *c*-subunits, explaining the cooperation observed in biochemical assays.

## Introduction

F_o_F_i_-ATP synthase (hereafter F_o_F_1_) is a ubiquitous enzyme that synthesizes or hydrolyzes ATP coupled with proton translocation at the inner mitochondrial membrane, chloroplast thylakoid membrane, and bacterial plasma membrane^1–3^. F_o_F_1_ synthesizes ATP via rotation of the central rotor driven by the proton motive force across the membrane. The enzyme comprises two rotary motors that share the rotor, i.e., the water soluble F_1_, which has catalytic sites for ATP synthesis/hydrolysis^4^, and the membrane-embedded F_o_, which mediates proton translocation^5^. The F_o_ motor consists of a *c* oligomer ring (*c*-ring), which serves as the rotor, and the *ab*_2_ stator portion located on the *c*-ring periphery. Downgradient proton translocation through F_o_ drives rotation of the central rotor composed of *c*-ring and γε subunits, thereby inducing conformational changes in F_1_ that result in ATP synthesis. Conversely, ATP hydrolysis in F_1_ induces reverse rotation of the rotor, which forces F_o_ to pump protons in the reverse direction.

The *c*-ring is composed of 8–17 *c*-subunits depending on the species^6–12^. F_o_F_1_ from thermophilic *Bacillus* PS3 and yeast mitochondrial F_o_F_1_ contain 10 *c*-subunits in the *c*-ring, which is designated the *c*_10_-ring^7,8,13,14^ (Fig. 1). The F_o_*-c* subunit harbors an essential proton-binding carboxyl group (*c*-Glu; cE56 in *Bacillus* PS3, *c*E59 in yeast mitochondria) located near the center of the membrane-embedded region; this group functions as the proton carrier (Fig. 1a). Protonation in the Glu allows the *c*_10_-ring to bind a proton, whereas proton release leads to Glu deprotonation. Accordingly, bacterial F_o_F_1_ activity is significantly decreased when the corresponding key residue is modified by the inhibitor. *N*,*N*-dieveloliewiearbodiimide (DCCD)^15^ or mutated to other amino acids^16^, and *Bacillus* PS3 F_o_F_1_ carrying a single *c*E56Q mutation in the *c*_10_-ring does not catalyze ATP-driven proton pumping or ATP synthesis^8^.

**Figure 1.**
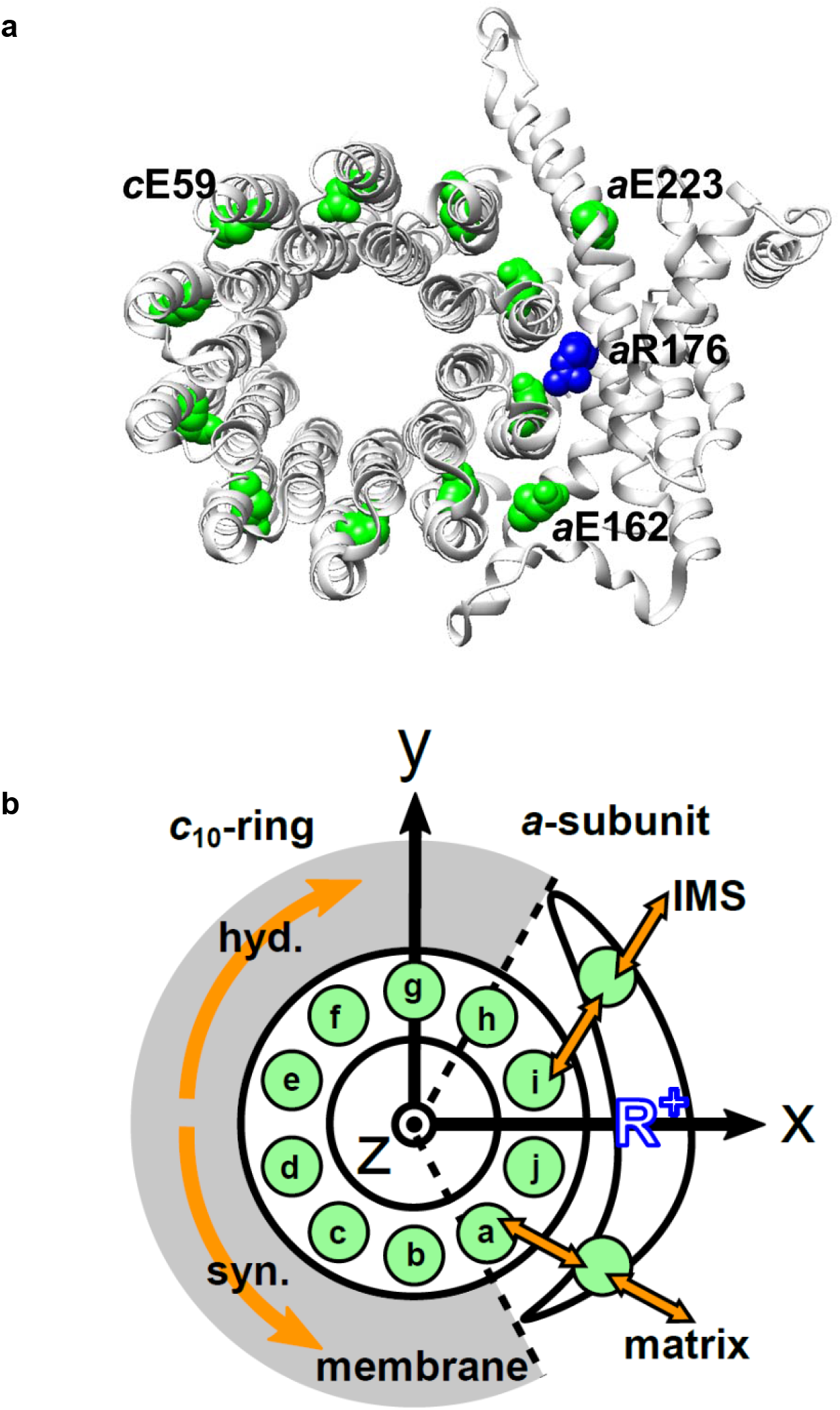
Schematic picture of the *a*-subunit and *c*-ring of F_o_. (**a**) The *ac*_10_ part of the F_o_ region is depicted as a ribbon diagram. Spheres represent *c*E59, which was substituted in this study, *a*E223, *a*E162, and *a*R176 (blue) (the residue numbers are those from yeast). (**b**) Schematic diagram of our simulation model. Green circles represent protonatable glutamates. Those in 10 *c*-subunits are labeled a-j. The membrane drawn in gray is modeled implicitly. Protons can hop between *c*E59 and glutamates in *a*-subunit, *a*E223, and *a*E162. Additionally, *a*E223 and *a*E162 exchange their protons with the IMS and matrix aqueous environment, respectively. Arrows in orange indicate the net proton flow. We set the rotational axis of the *c*_10_-ring as the z-axis, and the position of *a*R176 as the x-axis. Clockwise rotation of the *c*-ring occurs in ATP hydrolysis mode, and counterclockwise rotation of the *c*-ring occurs in ATP synthesis mode.

The *a*-subunit comprises two separate half-channels, one connecting the *c*-ring to the periplasm side of the bacteria or intermembrane space side of the mitochondria, while the other connecting the *c*-ring to the cytoplasmic side of the bacteria or matrix side of the mitochondria (Fig. 1b). Recent cryo-electron microscopy (EM) structural analyses of F_o_F_1_ at near atomic resolution^17,18^ have revealed two long tilted parallel α-helices in the *a*-subunit at the interface with the *c*_10_-ring. An essential Arg residue (*a*R169 in *Bacillus* PS3, *a*R176 in yeast mitochondria) at the middle of the long parallel helices plays a critical role in separating the two half-channels by preventing proton leakage^19^, and in the half-channels, two highly conserved Glu residues (*a*E223 and *a*E162 in yeast mitochondria) are regarded as proton-relaying sites^20^ (Fig. 1a). Since the essential Arg (*a*-Arg) localizes near *c*-Glu in the *c*_10_-ring, the attractive interaction between *a*-Arg and deprotonated *c*-Glu is hypothesized to also contribute to F_o_ rotation^21^.

In F_o_ rotation models proposed based on experimental studies^21–23^, the *c*-subunits facing the *a*-subunit perform three functions (proton release, electrostatic interaction with *a*-Arg, and proton uptake) depending on their positions relative to the *a*-subunit. A high-resolution structure of yeast mitochondrial F_o_F_1_ showed four of the 10 *c*-subunits facing the *a*-subunit^20^. Three key residues, i.e., *a*Glu162, *a*R173, and *a*Glu223, localize between the *c*-Glu residues of the four *c*-subunits, suggesting that the *c*-Glu residues of adjacent *c*-subunits could cooperate through the *a*-subunit residues. A more recent theoretical study using a hybrid Monte Carlo/molecular dynamics (MC/MD) simulation based on a high-resolution structure showed that there can be two or three deprotonated *c*-Glu residues facing the *a*-subunit concurrently^23^, suggesting potential synchronization; however, no experimental studies have confirmed this cooperation between *c*-subunits.

To directly investigate the cooperation among *c*-subunits in the *c*_10_-ring, we used a genetically fused single-chain *c*-ring and analyzed the function of *Bacillus* PS3 F_o_F_1_ carrying hetero *c*E56D mutations. Biochemical assays showed that the ATP synthesis activity was reduced, but not completely inhibited, by a single *c*E56D mutation and was further reduced by double *c*E56D mutations. Importantly, among five double mutants, the activity was decreased more as the distance between the two mutation sites increased. To clarify the molecular mechanisms, we performed proton transfer-coupled molecular dynamics (MD) simulations of F_o_, in which the mutations were mimicked, reproducing the characteristics of the biochemical experiment. From the analysis of the simulation trajectories, we found that prolonged duration times for proton uptake in the two mutated *c*-subunits can be shared. As the distance between the two mutation sites increases, the degree of time-sharing decreases. Taken together, these results reveal the functional coupling between neighboring *c*-subunits.

## Results

### Biochemical assays using F_o_F_1_s with a fused c-ring harboring hetero mutations

To investigate potential cooperation among *c*-subunits in the *c*_10_-ring rotation driven by proton translocation, we generated F_o_F_1_ mutants harboring a hetero-mutated *c*_10_-ring from thermophilic *Bacillus* PS3. We previously produced a fusion mutant, *c*_10_ F_o_F_1_, in which 10 copies of the F_o_-*c* subunit in the *c*_10_-ring were fused into a single polypeptide and demonstrated that *c*_10_ F_o_F_1_ was active in proton-coupled ATP hydrolysis/synthesis^8^. Starting from *c*_10_ F_o_F_1_, we generated six mutant F_o_F_1_s harboring one or two hetero *c*E56D-mutated *c*-subunits. The single mutant carries a *c*E56D mutation in the *c*(e)-subunit (designated as mutant “e”), whereas the five double mutants, “ef,” “eg,” “eh,” “ei,” and “ej,” harbor two *c*E56D mutations, each in the corresponding *c*-subunits (see Fig. 1b for labeling of *c*-subunits).

F_o_F_1_ mutants carrying one or two *c*E56D substitutions in the *c*_10_-ring were expressed in host *Escherichia coli* cell membranes at approximately one-tenth the level of wild-type (WT) F_o_F_1_. Western blotting with anti-*c*-subunit antibodies confirmed *c*-subunit expression in all mutants (Fig. 2a). First, ATP synthesis activity was measured using inverted membrane vesicles containing mutated F_o_F_1_s (Fig. 2b). The activity of mutant “e” was decreased to 24% ± 1.8% of that of F_o_F_1_ carrying only the fusion mutation. Moreover, among the five double mutants, “ef” showed the highest activity (19.7% ± 2.3% of that of *c*_10_ F_o_F_1_), and the activity decreased as the distance between the two mutations increased (“eg”: 18.3% ± 1.3%; “eh”: 15.6% ± 1.8%; “ei”: 15.0% ± 0.6%; “ej”: 14.0% ± 0.5%). Next, ATP-driven proton pump activity was measured as quenching of the fluorescence of 9-amino-6-chloro-2-methoxyacridine (ACMA) caused by proton influx into the inverted membranes (Fig. 2c). In the case of mutant “e,” fluorescence quenching was decreased to 36.0% ± 6.2% of that measured for the *c*_10_ fusion mutant lacking the *c*E56D mutation (Fig. 2d).

**Figure 2.**
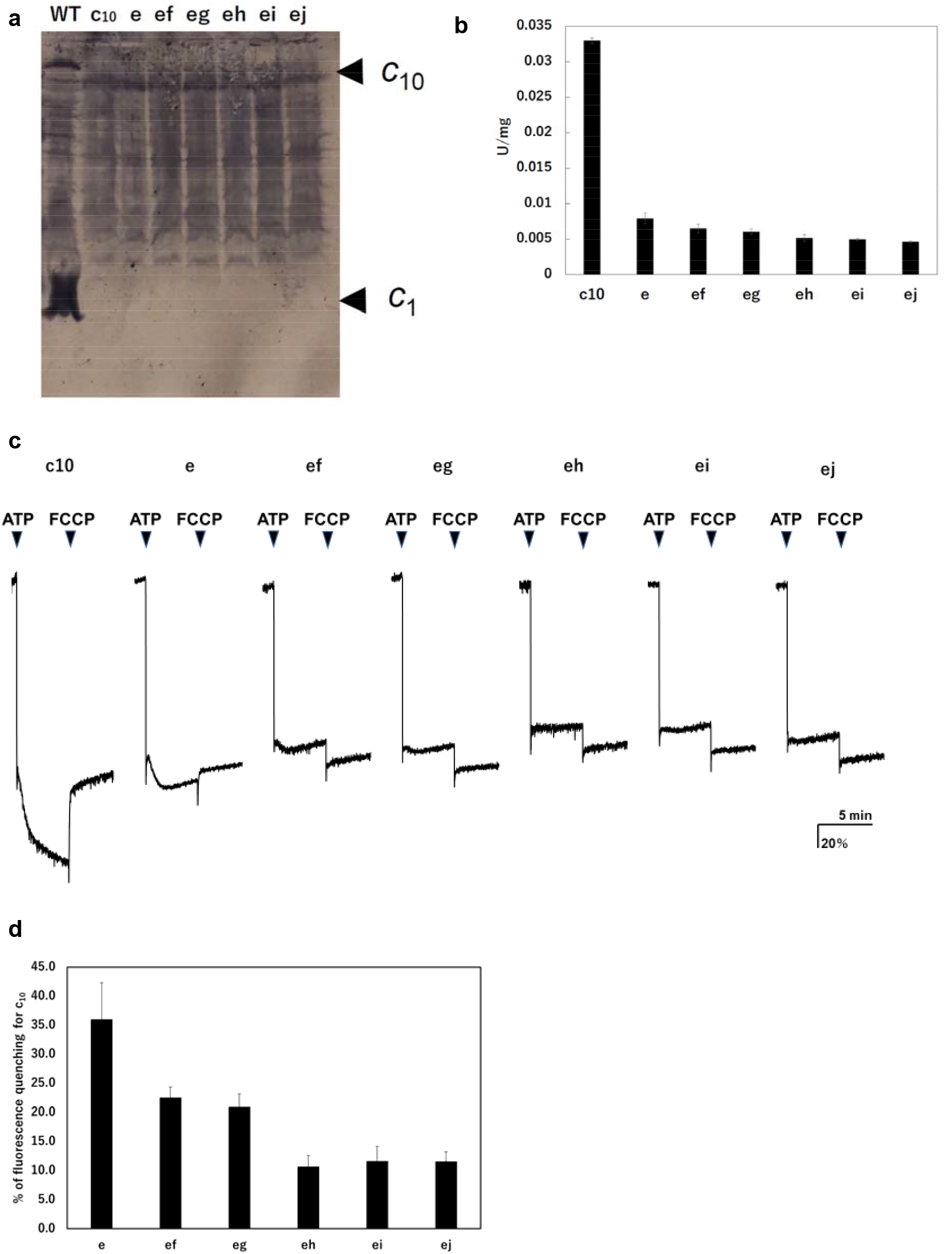
Expression of the mutated F_o_-*c* subunit and proton pump and ATP synthesis activities of membrane vesicles containing mutated F_o_F_1_s. (**a**) Proteins were separated using SDS-PAGE and immunoblotted with anti-F_o_-*c* antibodies. (**b**) ATP synthesis driven by NADH oxidation. (**c**) ATP-driven proton pump activity was measured by monitoring ACMA fluorescence quenching. (**d**) Percentage of fluorescence quenching for *c*_10_.

Among the double-mutants, “ef” and “eg” showed higher activity than the other mutants, with quenching to 22.6% ± 1.7% and 20.9% ± 2.2%, respectively, of that of *c*_10_ F_o_F_1_. The mutants “ei,” “eh,” and “ej” showed low fluorescence quenching; however, after the addition of *p*-trifluoromethoxyphenylhydrazone (FCCP), the fluorescence increased with a time constant of several seconds. This result indicated that protons were flowing into the inverted membrane vesicles and then flowed out following FCCP addition. The quenching ratios relative to *c*_10_ F_o_F_1_ calculated for mutants “ei,” “eh,” and “ej” were 10.7% ± 1.9%, 11.6% ± 2.5%, and 11.6% ± 1.6%, respectively. Thus, proton pump activity was high in the double mutants “ef” and “eg,” in which the two mutations are close to each other, but low in “eh,” “ei,” and “ej,” in which the mutations are introduced farther apart. Although the mutant F_o_F_1_s showed ATP hydrolysis activity, approximately 90% of the activity was insensitive to DCCD, a compound that inhibits F_o_ (Table 1). DCCD-insensitive ATP hydrolysis indicates uncoupled F_o_F_1_ activity. All mutants showed 10–15% DCCD-sensitive ATP hydrolysis activity. Thus, a subtle difference in the structure of the proton-binding site induced by the *c*E56D mutation may confer resistance to DCCD binding or cause uncoupling. The rotation driven by ATP hydrolysis is affected to a greater extent by the structure of the 3-fold symmetry of F_1_ than by the rotation during synthesis, and the DCCD-sensitive ATP hydrolysis activity indirectly reflects the function of F_o_.

**Table 1.** Membrane ATPase activities from cells expressing hetero-mutated *c*-subunits.

| Mutant | ATPase activity<sup>*</sup> |  |
| --- | --- | --- |
|  | –DCCD | +DCCD |
| *c*<sub>10</sub>-fusion | 0.15 | 0.067 |
| Mutant e | 0.087 | 0.076 |
| Mutant ef | 0.090 | 0.065 |
| Mutant eg | 0.078 | 0.070 |
| Mutant eh | 0.086 | 0.073 |
| Mutant ei | 0.088 | 0.080 |
| Mutant ej | 0.083 | 0.068 |

<sup>\*</sup>Membrane ATPase activity was measured after pre-incubation of membranes at 10 mg/mL in PA3 buffer with or without 50 μM DCCD for 20 min at 25 °C. Activity is expressed as μmol/min/mg.

### MD simulation of hetero-mutated F_o_F_1_s

Biochemical assays showed that the decreased rotation speed of the double-mutant F_o_ motor depends on the distance between the two mutation sites; however, the underlying mechanism was not clear. To obtain mechanistic insights, we tested the mutated F_o_ motor 23 rotations by proton transfer-coupled molecular simulations. Based on our previous simulation setup for the WT yeast mitochondrial F_o_, we introduced the *c*E59D mutation *in silico* to one and two *c*-subunits corresponding to the biochemical assays (see Methods for more details).

First, we demonstrated 10 trajectories for the single mutant “e” (Fig. 3a). Although the mutated *c*_10_-ring paused for a long period, the mutants still rotated in the synthesis direction coupled with proton transportation.

**Figure 3.**
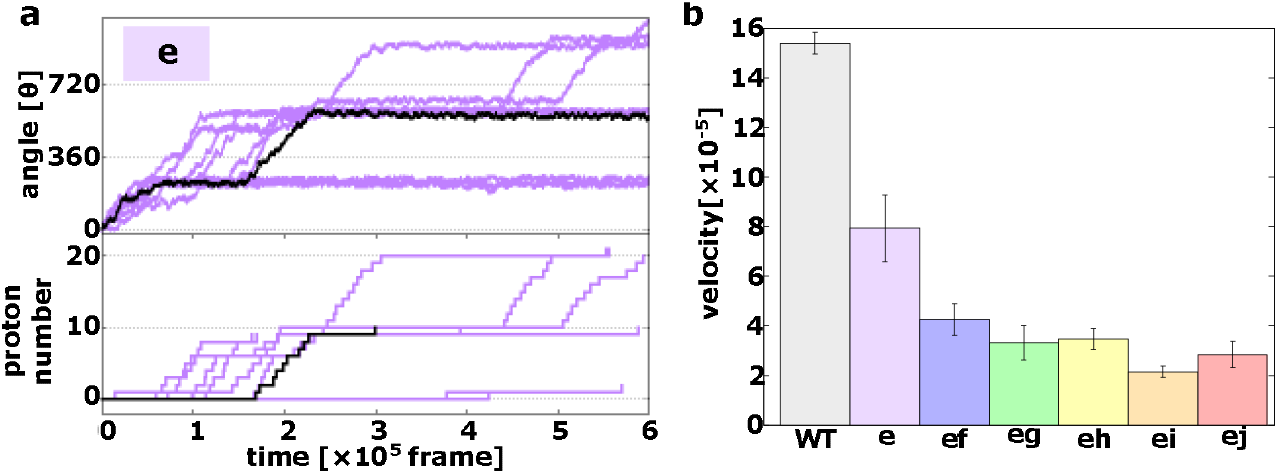
Proton transfer-coupled MD simulation of the WT and hetero mutants with Asp substitution of Glu. (**a**) Ten trajectories of the “e” mutant. The black line shows one representative trajectory. Upper part: rotation angle from initial position of *c*(a); lower part: the number of protons that entered from the IMS channel and were transported to the matrix channel through rotation. (**b**) Average rotational velocities for WT and mutants. Error bar: standard error.

Next, we simulated all five double mutants (“ef,” “eg,” “eh,” “ei,” and “ej”) and calculated the average rotational velocities over 10 trajectories (Fig. 3b). Fig. 3b shows the mean values and standard errors of the rotational velocities of the WT and all mutants. The rotational velocity of mutant “e” is almost two times slower than that of the WT. The rotational velocities of double mutants tend to decrease as the distance between the mutated chains increases. Thus, we were able to capture the characteristics of the experimental results in our simulations qualitatively, but not quantitatively.

We then evaluated the molecular processes for the simulation. Each *c*E59 (or *c*E59D) is protonated when the corresponding *c*-subunit is far from the *a*-subunit. This is regarded as the resting state of *c*E59 (Fig. 4a). As counterclockwise rotation occurs, the *c*-subunit approaches the half-channel of the *a*-subunit, which is connected to the matrix (the matrix half-channel). When *c*E59 comes close to *a*E162, which is the relaying site to the matrix half-channel, proton transfer from *c*E59 to *a*E162 occurs via the Monte Carlo step. Depending on the transfer efficiency, several Monte Carlo steps may be required to achieve proton release from *c*E59. We define the time from the first trial of the *c*E59-to-*a*E162 proton transfer to the success of transfer as “the duration for proton release” (pink in Fig. 4a). Once *c*E59 is deprotonated, the corresponding *c*-subunit can rotate counterclockwise further into the *a*-subunit facing region. After some rotation, the *c*-subunit approaches the other half-channel connected to the inner membrane space (IMS) (the IMS half-channel). When *c*E59 comes close to *a*E223, which is the relaying site for the IMS channel, *c*E59 attempts to take up a new proton from *a*E223 via the Monte Carlo step. We define the time from the success of proton release to the arrival at the rotation angle for proton uptake as “the duration for the deprotonated rotation” (indicated in green in Fig. 4a). Again, several Monte Carlo steps may be required to achieve this proton uptake. We define the time from the arrival at the proton uptake angle to success of proton uptake as the “the duration for proton uptake” (blue in Fig. 4a). Then, the *c*-subunit returns to the resting state. Thus, the entire time could be divided into three stages: stage 1, the duration for proton release; stage 2, the duration for deprotonated rotation; and stage 3, the duration for proton uptake, in addition to the resting time. Note that these durations are defined for each *c*-subunit and that the durations in one *c*-subunit overlap with durations in other *c*-subunits. For each mutant and for the WT, for each of the 10 *c*-subunits, we analyzed these three durations.

**Figure 4.**
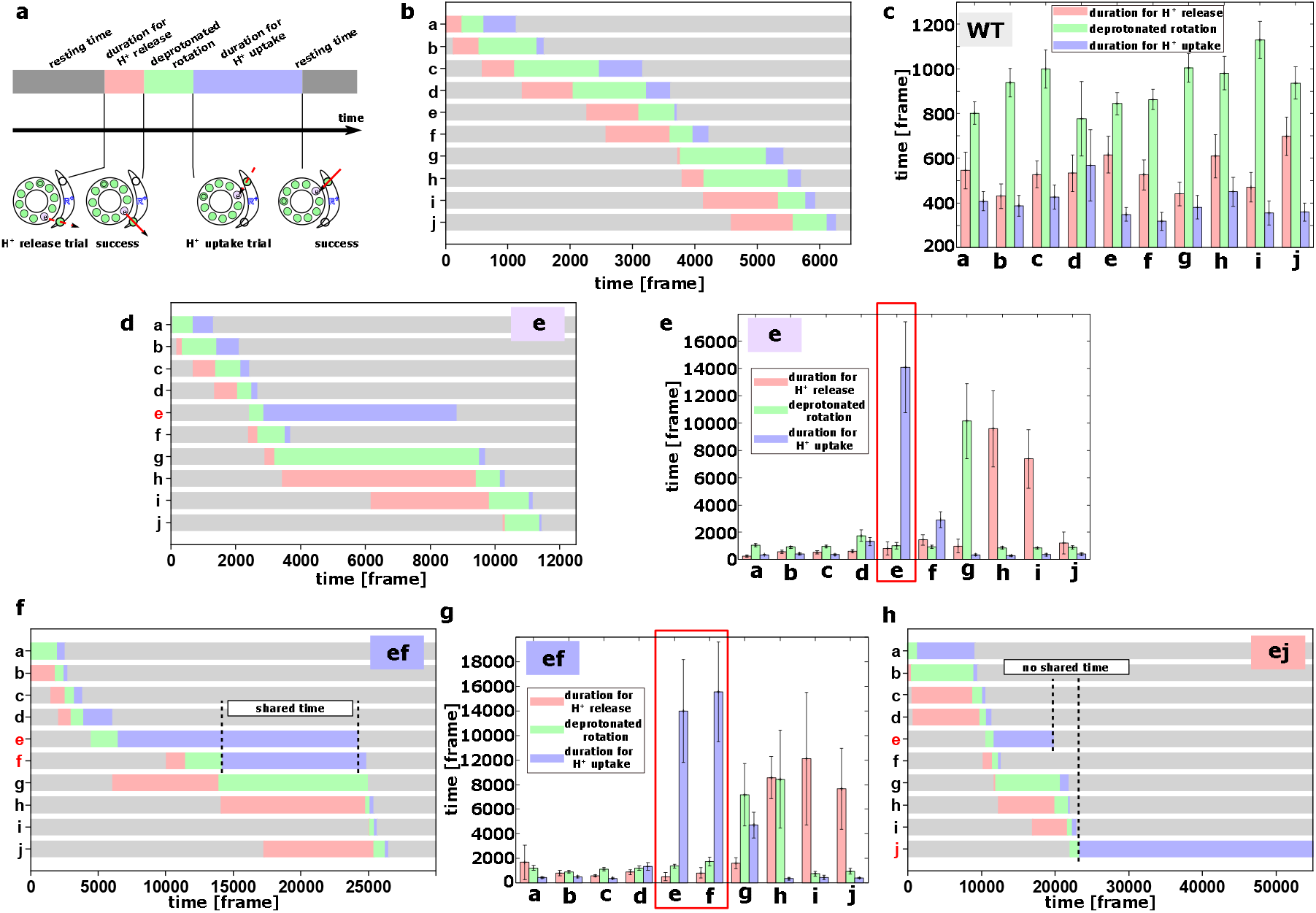
Analysis of the molecular simulations. (**a**) Schematic graph of duration times. The entire time was divided into the duration for proton release, the duration for deprotonated rotation, the duration for proton uptake, and the resting time. (**b**) Representative time course of durations for the WT. (**c**) Histogram of durations for every *c*-subunit of the WT. (**d**) Representative time course of durations for the single mutant “e.” (**e**) Histogram of durations for the single mutant “e.” (**f**) Representative time course of durations for the double-mutant “ef.” (**g**) Histogram of durations for the double-mutant “ef.” (**h**) Representative time course of durations for the double-mutant “ej.”

First, we examined the time course of a representative trajectory and the average durations for the WT for each *c*-subunit (Fig. 4b, c). The average durations for stages 1, 2, and 3 were approximately 500, 1100, and 200 MD frames, respectively. As expected, there were no significant differences in the durations among the 10 *c*-subunits.

Next, we performed the same analysis using the single mutant “e” (Fig. 4d, e). We found that the mutation in the *c*(e)-subunit clearly affects the duration for this subunit; stages 1 and 2 did not differ much from those in the WT, whereas the duration for stage 3 was much longer than that in the WT, as *c*E59D has a lower rate of proton transfer, and the pKa value of *c*E59D is lower than that of *c*E59.

We then analyzed double mutants. For the “ef” mutant (Fig. 4f, g), similar to the “e” mutant, the *c*E59D mutation in the *c*(e)-subunit prolongs the duration for proton uptake. Additionally, mutation in the *c*(f)-subunit prolongs the duration for proton uptake. Interestingly, as shown in Fig. 4f, these prolonged durations in *c*(e)- and *c*(f)-subunits are shared. Thus, by overlapping the delayed steps, the overall slowdown in the “ef” doublemutant system is lower than that if the effects of the two mutations were independent. In other words, sharing the delayed times of multiple subunits reduces the overall delay. In comparison, we examined the durations for the “ej” double-mutant (Fig. 4h).

As expected, mutations in the *c*(e)- and *c*(j)-subunits slow proton uptake in these subunits, although the durations are not shared. Therefore, we expect that there is no coupling between the *c*(e)- and *c*(j)-subunit mutations, resulting in additive effects of the two mutations.

In summary, coarse-grained MD simulations qualitatively reproduced the effects of single and double mutants found in biochemical assays and provided molecular interpretations of the coupling between two mutations. When the two mutations are in distant subunits of the *c*-ring, the effects of the two mutations are additive. In contrast, two mutations in neighboring subunits can result in overlapping of delays by the two mutations, leading to reduced effects of the two mutations.

### Simple kinetic analysis of hetero-mutant experiments

Although the biochemical data suggested some coupling among *c*-subunits, it remains unclear as to how they were coupled. MD simulation results suggested that cooperation can arise through sharing durations of proton release, deprotonated rotation, and proton uptake among a few *c*-subunits. Therefore, guided by the insight from the simulations, we next performed a simple kinetic analysis of the experimental data for the ATP synthesis rate (Fig. 2b).

We began with a simple case in which the 360° rotation is composed of a series of ten 36° rotation steps, with each step being independent. Then, for the WT, the rate constant *k_tot(WT)_* for 360° rotation can be expressed as

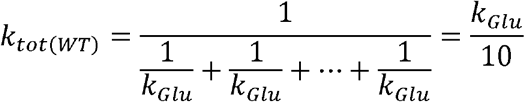

where *k_Glu_* is the rate constant for the WT *c*-subunit to rotate 36°. For the single mutant “e” bearing *c*E56D substitution at the *c*(e)-subunit, the rate constant *k_tot(e)_* for 360° becomes

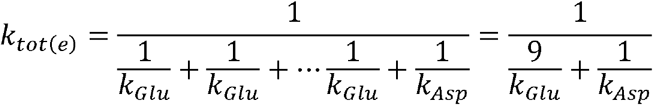

where *k_Asp_* is the rate constant for the *c*-subunit bearing the *c*E56D substitution to rotate 36°. From the measured ATP synthesis rates in Fig. 2b, we obtain

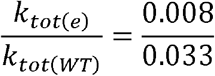

which, together with the above expressions, leads to

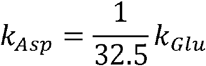

Using this relationship and the assumption of step independence, we can predict the rate constant of the mutually independent double-mutant as

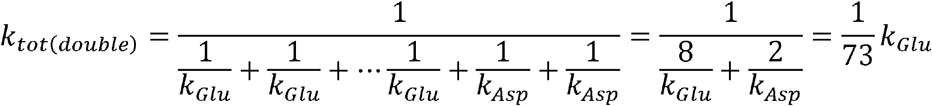

which, together with the expression for the WT, leads to

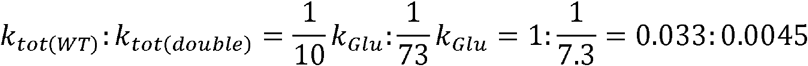

This rough estimate of *k_tot(double)_* ~0.0045 *U* is consistent with the ATP synthesis rate of the mutant “ej” (Fig. 2b). However, the rate of the mutant “ef” is clearly faster than the predicted value. This result suggests that clear separation of the two *c*-subunits (i.e., “ej”) shows no detectable coupling, whereas neighboring *c*-subunits (i.e., “ef”) show a negative coupling; that is, the effects of *c*E56D substitutions are reduced compared to the independent model.

Motivated by the suggestion from MD simulations that cooperation can arise by sharing the duration times among a few *c*-subunits, we then introduced the fraction of time shared between the mutated *c*-subunits as *x*. Then, the rate constant *k_tot(coupled)_* for 360° rotation can be expressed as

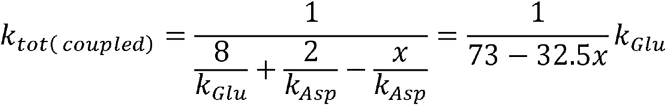

which leads to

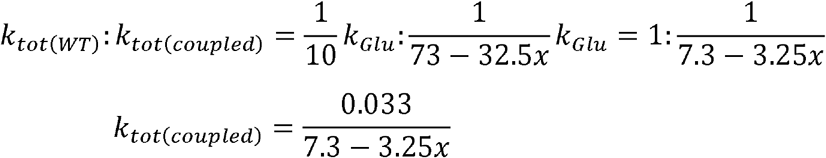

We can estimate the fraction of shared time of the hetero mutants “ef,” “eg,” “eh,” “ei,” and “ej” as 68%, 57%, 28%, 20%, and 5%, respectively. This simple analysis indicates that there is cooperation between two or more of the *c*-subunits and that the cooperation is weakened as the two subunits become further separated.

## Discussion

In this study, we determined whether *c*-subunits function in a cooperative manner for the rotation of the F_o_F_1_ *c*10-ring and assessed the mechanistic role of *c*-Glu (*c*E56) in this cooperativity. We have demonstrated that the degree of cooperation between two *c*-subunits depends on the distance between the *c*E56D hetero-mutation at the proton-carrying site. The activity of F_o_F_1_ was significantly decreased, but not completely abolished, by a single *c*E56D mutation. The activity was further decreased by the second *c*E56D mutation; moreover, the activity was high when the two mutations were introduced into nearby *c*-subunits, and the activity decreased as the distance between the two mutations increased. To the best of our knowledge, this is the first study providing unambiguous evidence for the coupling between two *c*-subunits. Molecular simulations reproduced the major features of biochemical experiments on single and double mutants and further revealed the molecular mechanisms of the coupling. Sharing of the prolonged durations by mutations in neighboring *c*-subunits leads to coupling.

When the *c*E56D substitution was introduced in one of the *c*-subunits, ATP synthesis activity was decreased substantially. In *E. coli* F_o_F_1_, ATP-driven proton pump activity was reported to be decreased after substitution of the conserved *c*Asp61 residue with Glu^24^. Here, after *c*E56D substitution, we detected partial retention of not only proton pump activity but also ATP synthesis activity. In contrast, *c*E56Q substitution in one of the *c*-subunits was found to eliminate ATP synthesis activity, ATP-driven proton pump activity, and DCCD-sensitive ATP hydrolysis activity^8^. In this study, ATP synthesis activity and ATP-driven proton pump activity were not completely lost when the carboxyl group of Glu was replaced with that of Asp. A comparison of this result with the *c*E56Q substitution results suggested that the presence of a carboxyl group capable of undergoing protonation and deprotonation is critical for rotation in the ATP synthesis direction coupled with proton transfer and for the proton-transfer-coupled rotation induced by ATP hydrolysis. As changing the Glu side chain to an Asp side chain decreased activity, we concluded that subtle structural differences in the proton-binding site caused by the one-methylene-group difference in the side-chain length, together with the change in pKa, slowed the elementary process required for driving rotation.

In F_o_F_1_s, carrying the *c*E56D mutation in two *c*-subunits, ATP synthase activity was high when the two introduced Asp residues were close to each other, and the activity decreased as the distance between the two mutations increased. If the kinetic bottleneck in the *c*_10_-ring rotation was only in one step of one *c*-subunit, the same activity would appear among double mutations with different relative separations. Alternatively, even if the *c*-subunit plays multiple roles, if each role works independently, the same activity would be obtained, irrespective of the mutational position. However, the experimental results showed that the activity was decreased when the two mutations were introduced farther apart. Thus, the data unambiguously indicate that the kinetic bottleneck in the *c*-ring rotation contains multiple *c*-subunits.

According to previously proposed models, proton release at *c*-Glu, electrostatic interaction between *a*-Arg and *c*-Glu, and proton binding at *c*-Glu drive *c*-ring rotation^21,22^. Moreover, based on the crystal structure of mitochondrial F_o_F_1_, the *c*-subunits that face the *a*-Glu223 residue bridging a proton from IMS, the *a*-Arg residue involved in electrostatic interaction, and the *a*-Glu162 residue bridging a proton to the matrix are located apart; therefore, we hypothesized that *c*-subunits on the *a*-Glu223 side of *a*-Arg play a role in proton release, whereas *c*-subunits on the *a*-Glu162 side of *a*-Arg play a role in proton uptake in the ATP synthesis rotation. MC/MD simulations based on the F_o_F_1_ atomic structure have revealed that proton transfer causes *c*_10_-ring rotation^23^. Here, MD simulation of *c*_10_-ring rotation during ATP synthesis was performed based on the aforementioned hypothesis that proton release and proton uptake are affected by the *c*E56D mutation. Our results indicated that the rotation speed is higher when the mutation is introduced at adjacent positions and that the rotation speed decreases as the distance between the two mutants increases. These results, which are consistent with the findings of our biochemical experiments, indicate cooperative proton uptake during the rotation of the *c*_10_-ring. Further analysis revealed that the waiting times for proton uptake in multiple subunits are shared. However, as the distance between the mutations increases, the degree of sharing of waiting time decreases, resulting in lower rotation speeds.

Overall, these findings suggested that at least three of the *c*-subunits on the *a*/*c* interface cooperate during *c*_10_-ring rotation in F_o_. This is consistent with the presence of two or three deprotonated carboxyl residues facing the *a*-subunit in the MC/MD simulation of WT F_o_F_1_^23^.

One limitation of this study is that we used the fusion mutation and the *c*E56D mutation. These mutations may affect not only our hypothesized driving force but also other activities. However, we consider our interpretations of the results to be valid based on the comparison with the combination of the same mutation and the results of MD simulations. Second, our MC/MD model includes only the *a*-subunit and *c*_10_-ring, whereas naturally occurring F_o_F_1_ also contains F_1_ and *b*-subunit. As F_1_ exhibits 3-fold symmetry, which is mismatched with the 10-fold symmetry in the *c*_10_-ring, the entire F_o_F_1_ is expected to exhibit more complex and asymmetric behaviors, which can represent a direction for future investigation of the enzyme.

## Methods

### Preparation of F_o_F_1_s carrying hetero mutations using fused multimeric F_o_-c

Plasmids for F_o_F_1_ mutants were generated from pTR19-ASDS^25^ using the megaprimer method and were then used for transformation of an F_o_-deficient *E. coli* strain, JJ001^26^. A plasmid for expressing the F_o_F_1_ mutant harboring a substitution of F_o_-*c* Glu-56 with Asp (*c*E56D) was prepared from pTR19-ASDS^25^ using the megaprimer method; this yielded pTR19-CE56D. The *c*E56D mutation sequence was verified through DNA sequencing. F_o_F_1_ carrying a hetero-mutation of *c*E56D in a fused *c*_10_-subunit prepared using Gly-Ser-Ala-Gly linkers^8^ was generated as follows. Briefly, an *AvrII* restriction site was introduced immediately after the initial *c*-subunit codon in the pTR19-CE56D expression plasmid, and new *NheI* and *SpeI* sites were introduced at downstream sites in the F_o_-*c* gene (to obtain pTR19-ACE56DN); pTR19-ACE56DN was digested with *Eco*RI and *Nhe*I, and the 1.3 kb *Eco*RI-*Nhe*I fragment was ligated into an *Eco*RI-*Avr*II site in pTR19-AC1N or pTR19-ACE56DN (to obtain pTR19-AC2DE or pTR19-AC2DD). Next, pTR19-AC2DE was digested with *Eco*RI and *Nhe*I, and the *EcoRI-NheI* fragment was ligated into an *Eco*RI-*Avr*II site in pTR19-AC1N or pTR19-ACE56DN (to obtain pTR19-AC3DEE or pTR19-AC3DED). By using this procedure, *uncE* genes were singly fused to generate plasmids expressing six F_o_F_1_s containing tandemly fused decamers carrying the *c*E56D mutation at the first hairpin (mutant “e”), first and second hairpins (“ef ‘), first and third hairpins (“eg”), first and fourth hairpins (“eh”), first and fifth hairpins (“ei”), and first and sixth hairpins (“ej”). The multimer *uncE* genes of the mutants were verified through plasmid restriction mapping. Plasmids generated for the WT and mutant F_o_F_1_s were singly expressed in F_o_-deficient *E. coli* strain JJ001 (*pyrE41*, *entA403*, *argHI*, *rspsL109*, *supE44*, *uncBEFH*, *recA56*, *srl*::*Tn10*)^26^. Transformants were cultured, and membrane vesicles were prepared as previously described^8^.

### Analytical procedures

ATPase activity was measured using an ATP-regenerating system at 37 °C in 50 mM Hepes-KOH buffer (pH 7.5), containing 100 mM KCl, 5 mM MgCl_2_, 1 mM ATP, 1 μg/mL FCCP, 2.5 mM KCN, 2.5 mM phosphoenolpyruvate, 100 μg/mL pyruvate kinase, 100 μg/mL lactate dehydrogenase, and 0.2 mM NADH^8^. One unit of activity was defined as hydrolysis of 1 μmol of ATP per minute; the slopes of decreasing 340 nm absorbance in the steady-state phase (400-600 s) were used for calculating activity. The sensitivity of ATP hydrolysis activity to DCCD-induced inactivation was analyzed as previously reported^4^. The ATP hydrolysis activity in the presence of 0.1% lauryldimethylamine oxide was measured to estimate the amount of F_o_F_1_ in the membrane vesicles. ATP-driven proton pump activity was measured as the fluorescence quenching of ACMA (excitation/emission: 410/480 nm) at 37 °C in 10 mM Hepes-KOH (pH 7.5), 100 mM KCl, and 5 mM MgCl_2_, supplemented with membrane vesicles (0.5 mg protein/mL) and ACMA (0.3 μg/mL)^8^. The reaction was initiated by adding 1 mM ATP, and quenching reached a steady level after 1 min; after 5 min, FCCP (1 μg/mL) was added, and fluorescence reversal was confirmed. The magnitude of fluorescence quenching at 3 min relative to the level after FCCP addition was recorded as the proton pump activity. ATP synthesis activity was measured at 37 °C using luciferase assays as previously described^27,28^. After incubating inverted membranes (5 mg/mL) with 5 mM N-ethylmaleimide for 15 min at room temperature, we added 1.6 mL PA3 buffer (10 mM Hepes-KOH [pH 7.5], 10% glycerol, and 5 mM MgCl_2_), 2.5 mM KPi (pH 7.5), 0.53 mM ADP (Calbiochem, San Diego, CA, USA), 26.6 μM P^1^,P^5^-di(adenosine-5□) pentaphosphate (Sigma-Aldrich, St. Louis, MO, USA), 20 μL inverted membranes, and 0.125 volumes CLS II solution (ATP Bioluminescence Assay Kit CLS II; Sigma-Aldrich) into the cuvettes; 0.5 mM NADH was added after starting the measurement. Synthesized ATP amounts were calibrated using a defined amount of ATP at the end of the measurement. Specific activity was calculated based on three parameters: estimated F_o_F_1_ concentration; slope of ATP synthesis activity measured for 50 s immediately after NADH addition (excluding slope of data recorded 100 s immediately before NADH addition); and ATP calibration value. FCCP addition was confirmed to prevent ATP synthesis. Protein concentrations were determined using a BCA assay kit (Thermo Fisher Scientific, Waltham, MA, USA), with bovine serum albumin serving as a standard. Membrane vesicles were separated using sodium dodecyl sulfate polyacrylamide gel electrophoresis (SDS-PAGE) with 15% gels containing 0.1% SDS, and proteins were stained with Coomassie Brilliant Blue R-250. F_o_F_1_ expression was confirmed by immunoblotting with anti-β and anti-*c* polyclonal antibodies for F_o_F_1_ from the thermophilic *Bacillus* PS3.

### Basic simulation system

To represent the proton transfer-coupled rotational motion of the *c*_10_-ring, protein motion and proton jump were modeled using MD and MC, respectively, and these dynamics were combined to reproduce *c*_10_-ring rotational motion with proton hopping^23^. In our simulation system, we included the *a*-subunit and *c*_10_-ring (Fig. 1a) structure models of yeast F_o_ based on the cryo-EM structure of a yeast mitochondrial ATP synthase (PDB ID: 6CP6)^20^. We used the AICG2+ coarse-grained model, where each amino acid is represented as a single particle located at the corresponding *Cα* atom. While lipids were not explicitly modeled, interactions between protein residues and the lipid membrane were represented through implicit membrane potential. Water solvents were also treated implicitly. The hybrid MC/MD simulations consisted of the MC phase, at which protonation states of 12 protonatable sites (the glutamic acid [or aspartic acid in the case of mutants] in 10 *c*-subunits, *a*E223, and *a*E162) are updated, and the MD phase, when amino acid positions are updated by Langevin dynamics. Each round contained MC trial moves for all the protons involved, followed by 10^5^ MD steps. All simulation setups were the same as those we have recently reported^23^, except for the treatment of the *c*E59D mutation.

### Treatment of cE59D in the simulation

In the hybrid MC/MD simulation, we mimicked *c*E59D mutations in the following manner. In the MD part, we simply changed the amino acid identity of the corresponding residue from glutamic acid to aspartic acid using the mutagenesis feature of PyMol. Given the nature of our coarse-grained representation, this results in minor changes. The MC move represents proton transfer, which must be largely affected by the *c*E59D mutations via two distinct mechanisms, i.e., the change in transfer efficiency and the change in the free energy difference between protonated and deprotonated states. For the former, the proton transfer efficiency is markedly reduced by the *c*E59D mutation because aspartic acid has a shorter sidechain than glutamic acid by one methylene-group. In our model, the transfer efficiency contains exp(—A(r — r_0_)) factor, where r is the distance between Cα atoms of the donor and the acceptor, the offset distance r_0_ represents the sum of sidechain lengths of the donor and acceptor, and *A* is the decay rate. We used r_0_ = 0.8 *nm* for *c*E59 (the same value as reported previously^23^) and set *r_0_* — 0.6 *nm* for *c*E59D, representing its shorter sidechain of aspartic acid. The decay rate *A* was set to 2.5 (1/nm) for *c*E59 (the same value as reported previously^23^) and 9.0 (1/nm) for *c*E56D. Second, the free energy difference between the states before and after the proton transfer is modulated by pKa differences in the donor and the acceptor amino acids and thus is affected by the *c*E59D mutation. Although the pKa value specific to the corresponding site is unknown, we empirically chose pKa = 8.0 for *c*E59 and 7.0 for *c*E59D considering the intrinsic difference in pKa values.

### Simulations and their analyses

For each of the WT F_o_ *ac*_10_ and the six *c*E59D mutation patterns corresponding to the biochemical assay, we carried out 10 independent simulation runs with different stochastic forces. The mutants included the single mutant “e” and the five double-mutants “ef,” “eg,” “eh,” “ei,” and “ej”. The single mutant “e,” for example, has the *c*E59D substitution only in the “e” chain, whereas other chains contain the WT *c*-subunit sequence. The double-mutant “ef” harbors substitutions in the two neighboring subunits. Each simulation run contained 6,000 rounds of MC/MD cycles (twice as long as in our previous paper^23^). Each round contained MC trial moves for all the protons involved and 10^5^ MD steps. Thus, the entire trajectory corresponds to 6.0 × 10^8^ MD (60,000 frames saved).

Notably, due to limitations in the computation time, we could simulate only one to a few turns of 360° rotations for each trajectory. As the mutant systems show asymmetric arrangements, the unbiased estimate of average velocities requires the rotation of multiples of 360°. Thus, we used the cumulative rotation angle and the MD time step at which the *c*_10_-ring returned to the initial orientation for the last time in each trajectory. The rotation velocity was obtained as the ratio of the cumulative rotation angle to the MD time. This velocity was then averaged over 10 trajectories.

## Acknowledgments

We thank Dr. Toshiharu Suzuki and Dr. Masasuke Yoshida for providing us with the expression plasmid for WT F_o_F_1_. This work was supported partly by a Grant-in-Aid for Scientific Research (C) and (B) [17K07922, 19H02577] and the Cooperative Research Program of “NJRC Mater. & Dev.”

## Author Contributions

N.M., S.K., and S.T. designed the research. N.M., S.O., H.T., and Y.S. performed experiments and analyzed the data. S.K. and T.N. developed the simulation code, performed simulations, and analyzed the data. N.M., S.K., and S.T. wrote the paper.

